## Supplemental Information for "The role of heterochronic gene expression and regulatory architecture in early developmental divergence"

*GO Enrichment Results*

Spreadsheet of all GO enrichment test results for various categories of relevant genes: https://github.com/NathanDHarry/Harry-Zakas-TC

*Supplementary figures and tables*

**SFigure 1**. Read mapping rate between samples. Mapping rates in all pairwise comparisons do not differ significantly. P-values are shown above the plot indicating no significant differences between the mapping rates of any two groups.

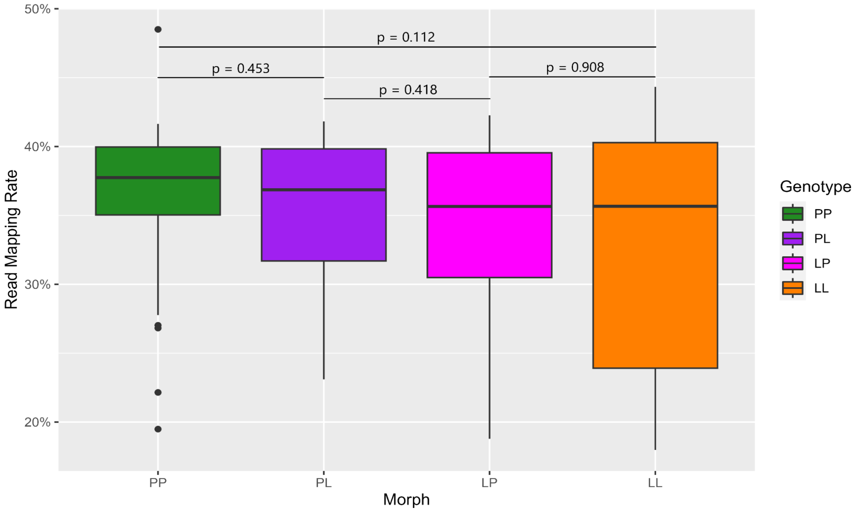


**SFigure 2**. Number of genes expressed at each stage in PP and LL samples.

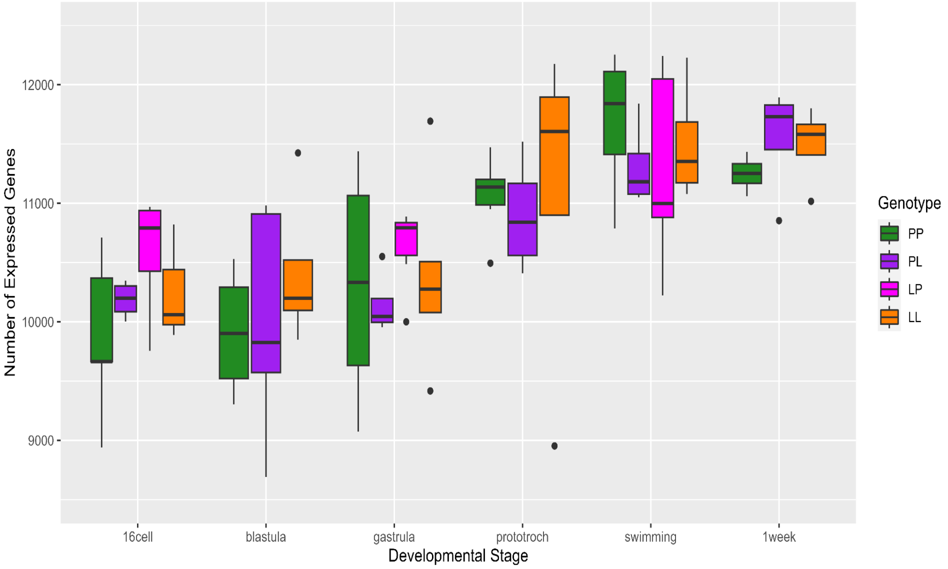

**Sfigure 3.** Cluster number optimization. The correlation between the two most similar gene expression clusters (profiles) as cluster number increases. Threshold (solid line) at r = 0.85.

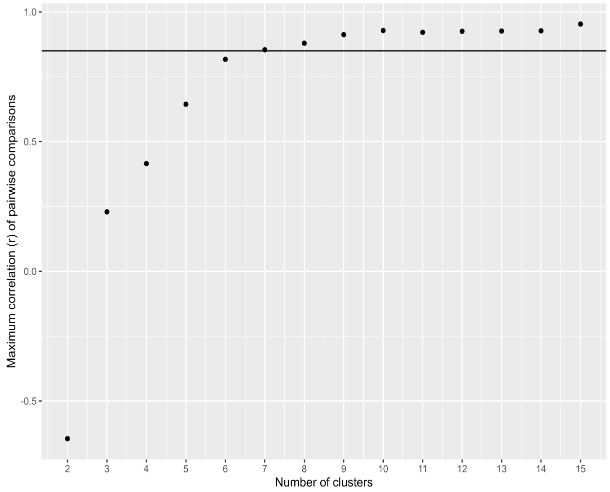


**Sfigure 4.** Clusters with all genes shown plotted on top of cluster core genes. Red lines indicate genes with high cluster membership scores while blue lines indicate genes with lower membership scores. Genes with no cluster membership score greater than 0.5 were not assigned to a cluster and are not presented in this study's results.

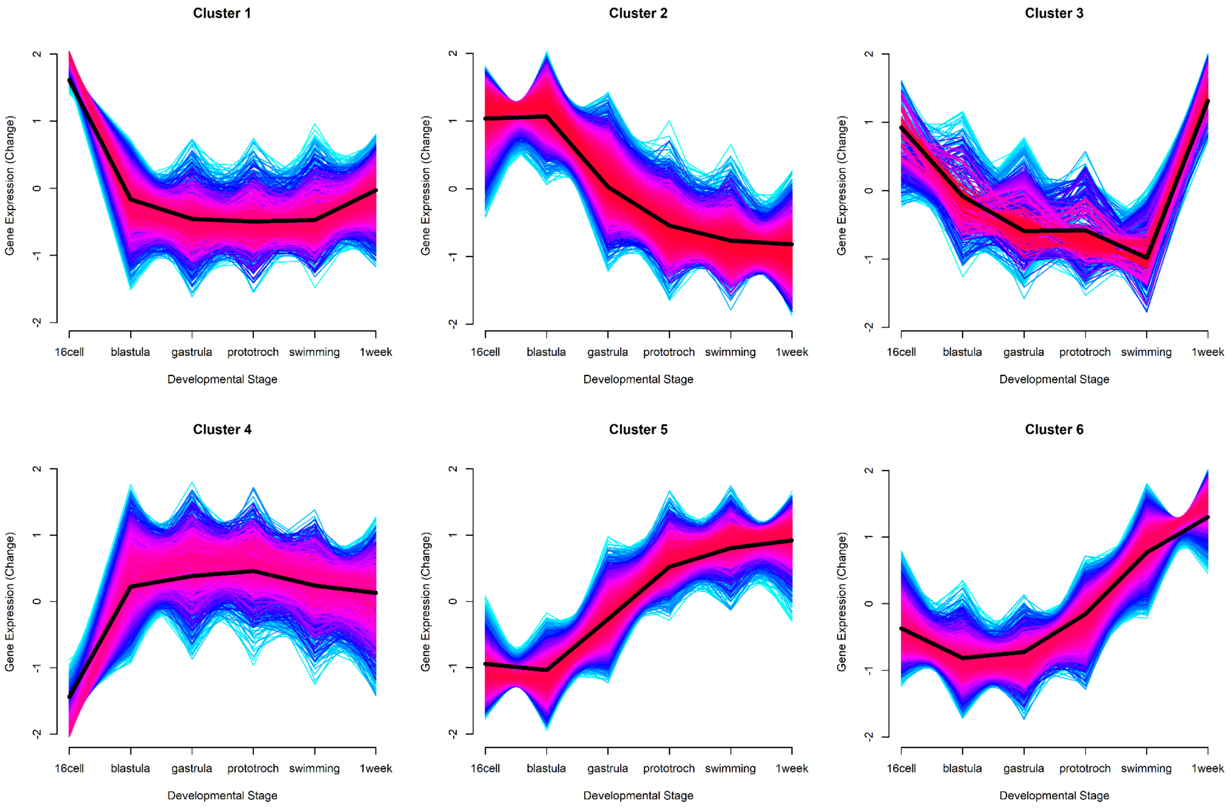

**SFigure 5.** Genes with parental effects. Most genes match maternal expression patterns.

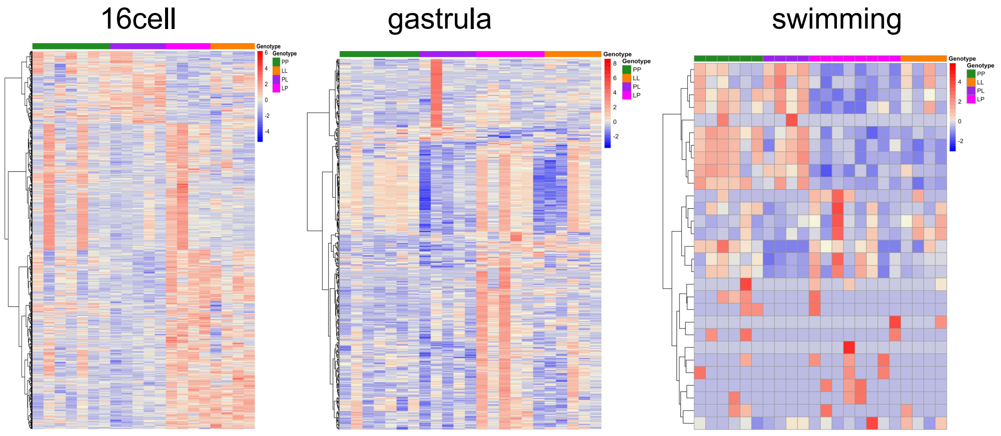


**SFigure 6.** PCA of all gene expression labeled by family (crossID).

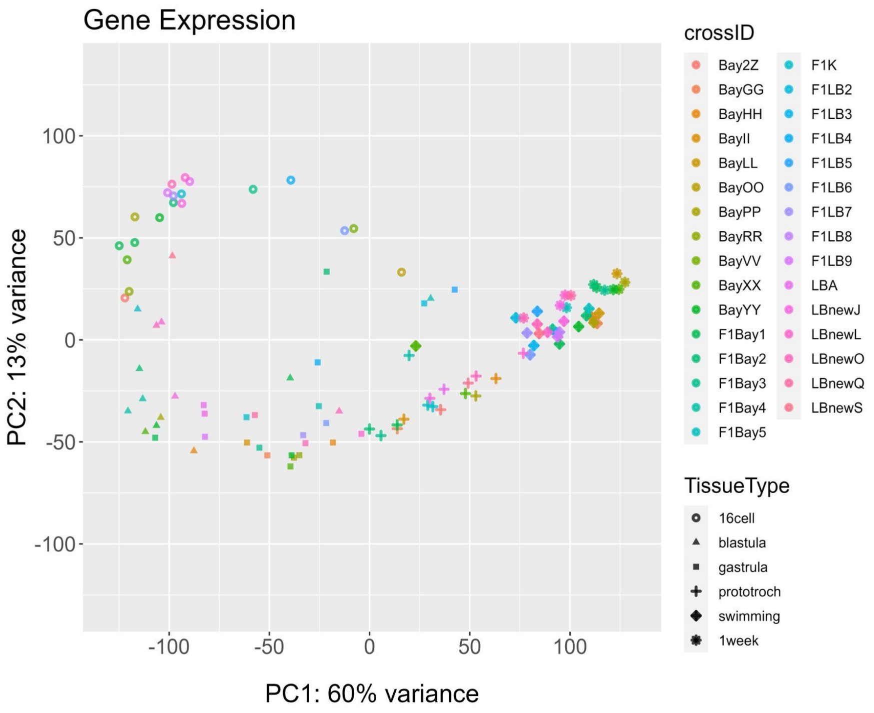


**STable 1**.

Mode of regulatory change classifications

| PP vs LL | $F_1$ P allele vs L allele | Ratio of DE | Classification |
|---|---|---|---|
| **PP ≠ LL** | $P_{allele} \neq L_{allele}$ | $PP/LL = P_{allele}/L_{allele}$ | Cis |
| **PP ≠ LL** | $P_{allele} = L_{allele}$ | $PP/LL \neq P_{allele}/L_{allele}$ | Trans |
| **PP ≠ LL** | $P_{allele} \neq L_{allele}$ | $PP/LL > P_{allele}/L_{allele}$ | Cis + Trans |
| **PP ≠ LL** | $P_{allele} \neq L_{allele}$ | $PP/LL < P_{allele}/L_{allele}$ | Cis x Trans |
| **PP = LL** | $P_{allele} \neq L_{allele}$ | $PP/LL \neq P_{allele}/L_{allele}$ | Compensatory |
